# Molecular dynamics simulations and linear response theories jointly describe biphasic responses of myoglobin relaxation and reveal evolutionarily conserved frequent communicators

**DOI:** 10.1101/682195

**Authors:** Bang-Chieh Huang, Lee-Wei Yang

## Abstract

In this study, we provide a time-dependent (td-) mechanical model, taking advantage of molecular dynamics (MD) simulations, quasiharmonic analysis of MD trajectories and td-linear response theories (td-LRT) to describe vibrational energy redistribution within the protein matrix. The theoretical description explains the observed biphasic responses of specific residues in myoglobin to CO-photolysis and photoexcitation on heme. The fast responses are found triggered by impulsive forces and propagated mainly by principal modes <40 cm^-1^. The predicted fast responses for individual atoms are then used to study signal propagation within protein matrix and signals are found to propagate ∼ 8 times faster across helices (4076 m/s) than within the helices, suggesting the importance of tertiary packing in proteins’ sensitivity to external perturbations. We further develop a method to integrate multiple intramolecular signal pathways and discover frequent “communicators”. These communicators are found evolutionarily conserved including those distant from the heme.

## Introduction

Understanding the allosteric control of molecular functions through investigations of energy flows within biomolecules has been of continuously arising scientific and medicinal interest in the past 20 years, encouraged by ever increasing computing power and implementations of appropriate physical theories. Exemplary systems include the relaxation dynamics of photolyzed heme-proteins[1-8] such as the photodissociation of monoxide (CO) originally captured in carbonmonoxy myoglobin (MbCO). This could be carefully studied by ultrafast spectroscopy that empowers scientist to characterize relaxation dynamics on the time scale of femto- to picoseconds[2, 3, 9, 10]. The energy flows of multiple stages relaxations started from the photoexcited electronic state of heme by ultrafast lasers, and then the energy flows through the vibrational excitation of heme to the proteins environment, and finally dissipate to the solvent.[1] Using the time-resolved near-IR absorbance spectra[1], it has been reported that the photoexcited electronic state of heme relaxes to its ground state with a time constant of 3.4 ± 0.4 ps and the ensuing thermal relaxation characterized by the temperature cooling exponentially with a time constant of 6.2 +- 0.5 ps.[1] In the crystalline environment, such a relaxation was reported to approach equilibrium at a faster pace through damped oscillations with a time period of 3.6 ps [11]. Further, the excited heme communicates with distant sites through vibrational modes within a few picoseconds, where the delocalized “doming modes” of heme are identified around 40 *cm*^−1^ using the nuclear resonance vibrational spectroscopy.[7] Two in-plane heme modes *ν*_4_ and *ν*_7_ are referred to couple with motions along the doming coordinate (40-50 *cm*^−1^) and with spatially extended modes (centered at 25 *cm*^−1^) by using high frequency laser pulses.[8]

Recently, the advancement of ultraviolet resonance Raman (UVRR) has been exploited to investigate the vibrational redistribution in protein matrix with the aid of its enhanced intensity of the Raman scattering.[2, 3, 9, 10] Mizutani’s group used time-resolved UVRR (UV-TR3) to measure time constants of relaxation dynamics for several tryptophan mutants in two different experiments of the photodissociation of MbCO.^[2, 3]^ In 2007, the CO photodissociation coupled with a conformational changes of Mb was studied. The band intensity change as a function of time owing to the hydrophobicity change upon the structural change in E helix as well as its interaction with A helix in which Trp7 and Trp14 are situated.^[3]^ The study also reported a biphasic changes in Trp14 and Tyr46 with a fast decaying response in a couple of picoseconds, followed by a slower recovery responses for 7 to >40 ps, involving permanent intensity changes.^[3]^ In 2014, the same group irradiated and excited the heme, and vibrational relaxation of Mb was monitored by UVRR under the condition that conformational change is suppressed where Fe is in the 3^+^ state.^[2]^ In these studies,^[2, 3]^ the biphasic decay of relaxation motions were observed and time constants of fast and slow responses of several residues have also been identified, which provides us an excellent model system to trace the energy flows and investigate the underlying mechanism of vibrational energy transfer within Mb.

On the theoretical side, many theories and algorithms have been developed to describe the vibrational energy relaxation and mechanical signal propagation.[12-18] The lifetime of CO vibrations estimated by the Landau–Teller formula is found to agree well with the time-resolved mid-IR absorbance experiments.[4] To further understand the inherent inhomogeneity in the spatially dependent relaxation rate of the solvated protein, Langevin model has been used to estimate inherent friction in protein motion.[19-21] Besides, the propagation of heat flow, kinetic energy or vibrational energy redistribution in proteins has also been investigated with mode diffusion (or mode-coupling) induced by anharmonicity,[22-25] MD simulations[26-28] and linear response theory (LRT).[21, 29-31]

In our previous work, we were able to use the normal mode based linear response theories (NMA-td-LRT)[21] to describe the relaxation dynamics of ligand photodissociation of MbCO in the UV-TR3 experiment as solvent-damped harmonic oscillators solved by Langevin dynamics.[3] A general assumption of the normal mode based theory is that the conformational space of protein motions is around the global minima of potential energy surface (PES). However, protein evolves across wider range of PES[32, 33] at the room temperature and the span of the motion can be described by PC modes derived from the essential dynamics or principal component analysis (PCA). Although this entails the concern of external perturbation that is no longer small, which challenges the validity of the td-LRT, we have earlier proved that, for systems evolving on a harmonic energy surface, the assumption of small perturbation is no longer a requirement [21, 34] (see also SI).

In this study, we formulate the PCA-based linear response theory (PCA-td-LRT) to investigate the signal propagation in myoglobin. The theoretical formulation is examined by comparing the estimated time constants of several residues for short (the “fast” response in the UVRR experiment) and long time (the “slow” response in the UVRR experiment) relaxation with two UV-TR3 experiments.^[2, 3]^ We also show that the large difference in energy-transfer speed for inter- and intra-helical signals, suggesting the importance of protein tertiary packing in mediating vibrational energy. We also explore the mode dependence of the signal propagation to understand the range of modes heavily involved in transmitting fast response signals. Finally, following a recently introduced communication matrix strategy to record the counts of mechanical signals[17] launched from sites nearest to the FE atom propagating through donor-acceptor pairs; the communication score (CS) of each residue is assigned, which quantifies how frequently mechanical signals goes through this residue shall perturbations are introduced in specific sites, such as substrate binding sites. The sites with high CS are termed “intramolecular communication centers (CCs)”, which are found in this study to be evolutionarily conserved. Also, these CCs were previously found able to allosterically regulate the enzyme activity[17].

## Methods

### MD protocols and Principal Component Analysis (PCA)

Unligated (1A6N) and CO-ligated (1A6G) myoglobin (Mb) structures are used in the MD simulations using the CHARMM36 forcefield implemented in NAMD package.[35] Proteins are immersed in a box of size 78Å×71Å×71Å containing 33276 TIP3P water molecules and neutralizing ions containing one sodium and nine chloride ions. Structures are energy-minimized, heated and equilibrated at 300K and 1 bar, maintained by Langevin thermostat and barostat. Particle Mesh Ewald and SHAKE are applied in the simulations. The production run at 300K and 1 bar takes 80 ns with snapshots being saved for every 100 fs. We take the non-hydrogen atoms of the last 50 ns, totaling 500,000 snapshots (frames), for the subsequent principal component analysis (PCA[36]). The structural change under perturbations 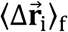 (see below) is taken from the difference between averaged ligated and unligated frames. The time-dependent and independent covariance matrices are calculated by PCA-derived quantities using the averaged unligated structure, to which the “0” of “< >_0_” in the eqs. (1) and (2) refer.

We performed (atom-)mass-weighted superimposition for each snapshot[37] of the unligated myoglobin onto the averaged structure in an iterative fashion.[36, 38] According to our earlier protocols,[36] the mass-weighted protein atom coordinates in an ensemble of superimposed frames were used to build the residue-residue covariance matrix for positional deviations, to which PCA was applied. PCA provided a set of principal component modes (PC modes) and their corresponding mass-weighted variance *σ*^2^ (the eigenvalues). According to equipartition theorem of harmonic oscillators,[36] at a given temperature T, the energy of a harmonic PC mode is *σ*^2^ × *ω*^2^ = *k*_*B*_*T*, where the effective frequency, ω, of the PC mode can be defined as 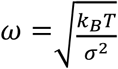.

### PCA-based time-dependent linear response theory

In the studies of photodissociation of carbonmonoxy myoglobin (MbCO),[2, 3] a biphasic relaxation - a fast response following by a long-time relaxation – for UV-TR3 -detectable residues was observed. To model the slow and fast (biphasic) relaxation dynamics corresponding to the residue responses with and without conformational changes,[2, 3] we developed two PCA-based time-dependent linear response theories (td-LRT)[21]- the constant force td-LRT (CF-td-LRT) and the impulse force td-LRT (IF-td-LRT), respectively. The CF-td-LRT reads as

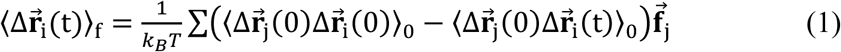

where *k*_*B*_ and *T* are the Boltzmann constant and temperature, respectively. The time dependent covariance matrix, 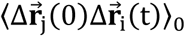, can be expressed as the sum of solvent-damped harmonic oscillators treated by the Langevin equation (see supporting information, SI). When time goes to infinite, 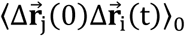 vanishes and eq. (1) retreats back to the time independent form,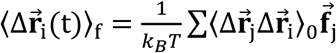.[31] The constant forces which drive Mb to evolve from unbound to bound conformation, upon CO binding, can be derived from the time independent LRT as the form 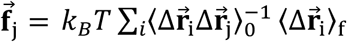 in kcal/mol/Å, where 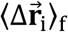 in this study is the structural difference between bound and unbound Mb structures that are the averaged MD snapshots. Here the initial structures for MD simulations were taken from the x-ray-resolved unligated (PDB:1A6N) and ligated Mb (PDB:1A6G) structures.^[21]^

To model the relaxation dynamics for UV-TR3 experiment without conformational changes, we use the IF-td-LRT^[21]^ in the form

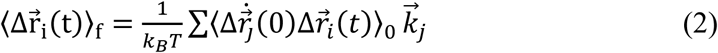

where 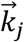 is an impulse force applied on the atom *j*. In this study, we model the laser pump pulse which deposits excess energy on heme by the force pointing from CO (if bound) to the FE atom plus a set of forces that model the “heme breathing” (*ν*_7_) mode.[8]

In both td-LRTs, we express the time-dependent covariance matrix with the PC modes subjecting to the Langevin damping[39] where the solvent friction, β of 27 cm^-1^ is taken from our earlier calculation[21] based on Haywards’ approach.[20] Under this friction constant, PC modes of frequency higher than 13.5 *cm*^−1^ are underdamped, which agrees with the observation of orientation-sensitive terahertz near-field microscopy that vibrational modes of frequency larger than 10 *cm*^−1^ are overdamped in a chicken egg white lysosome of 129 residues.^[40]^

**The mean characteristic time of a residue**,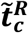

Using the IF-td-LRT in the Eq. 2, after an impulse force 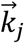 is applied on a *C*_*α*_ atom, atoms evolve as a function of time. After perturbations are introduced to the system at time 0, the atom *i* achieves its maximal displacement from the initial position at time *t*^*i*^. The time when 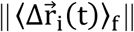 is at the maximum of all time is defined as the characteristic time of atom 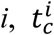. Our previous study[21] showed that characteristic time is a function of the directions of applied impulse forces. When a single point force is applied on one atom in the system, the 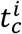 does not change with the absolute magnitude of the force and remains unchanged if the force points to the opposite direction. To average out the effects of applied force directions, we define a set 21 impulse forces, with unit magnitude and pointing toward evenly distributed 21 directions in a hemisphere at spherical angles of Ω, acting on a heavy atom of an interested residue. Consequently, a set of characteristic times 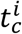 could be derived. We then averaged 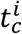 over 21 directions as well as all heavy atoms *i* of a residue *R*; the averaged characteristic time of a residue *R* is expressed as 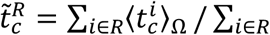 where *i* is the index of heavy atoms. To estimate the signal propagation speed, 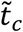 can be plotted as a function of the distance, 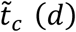, where *d* is the distance between the perturbed *C*_*α*_ atom and the *C*_*α*_ atom that senses the coming signal. By taking the linear regression for a set of 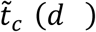 as a function of *d*, we can obtain the propagation speed as the reciprocal of the slope (see Figure 3).

### Communication matrix/map (CM) and communication score (CS)

On the aid of the IF-td-LRT providing the characteristic time 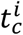 to characterize the dynamical signal propagation, we further develop a method that trace the signal propagation pathways and identify which residues are essential to mediate the signals when considering all the pathways. Here, we introduce “the communication map (CM)” to record the signal propagations between residues, from which we derive “the communication centers (CCs)” - the residues that are frequently used to communicate signals in most of the pathways. 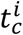 can provide causality of signal propagation. We could trace the signal transmission pathways consisting of the donor-acceptor pairs illustrated in Fig. 1. There are two communication criteria to determining the atom communication between a donor atom, *a*_*d*_, and an acceptor atom, *a*_*r*_. First, given an impulse force exerted on an atom in the system, the atoms with characteristic times that differ by Δ*t*, are considered “communicating” such that 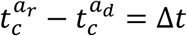, where the fixed Δ*t* is chosen to be 0.2 ps in this study. In order to maintain the causality of signal propagation, the candidates that satisfy the first criterion are required to meet the second criterion such that the angle between the vector pointing from the perturbed atom to *a*_*r*_ and that pointing from the perturbed atom to *a*_*d*_ should be less than 90 degrees. It is possible that a donor atom connects multiple acceptor atoms, or multiple donors connect to a single acceptor atom. Consequently, in order to quantitatively recognize how frequently a residue participates in the signal propagation, we define an all-atom communication matrix *F*(*a*_*d*_, *a*_*r*_) which account for the number of the communication that the donor, *a*_*d*_, connects to the acceptor, *a*_*r*_. As the form

**Figure 1.**
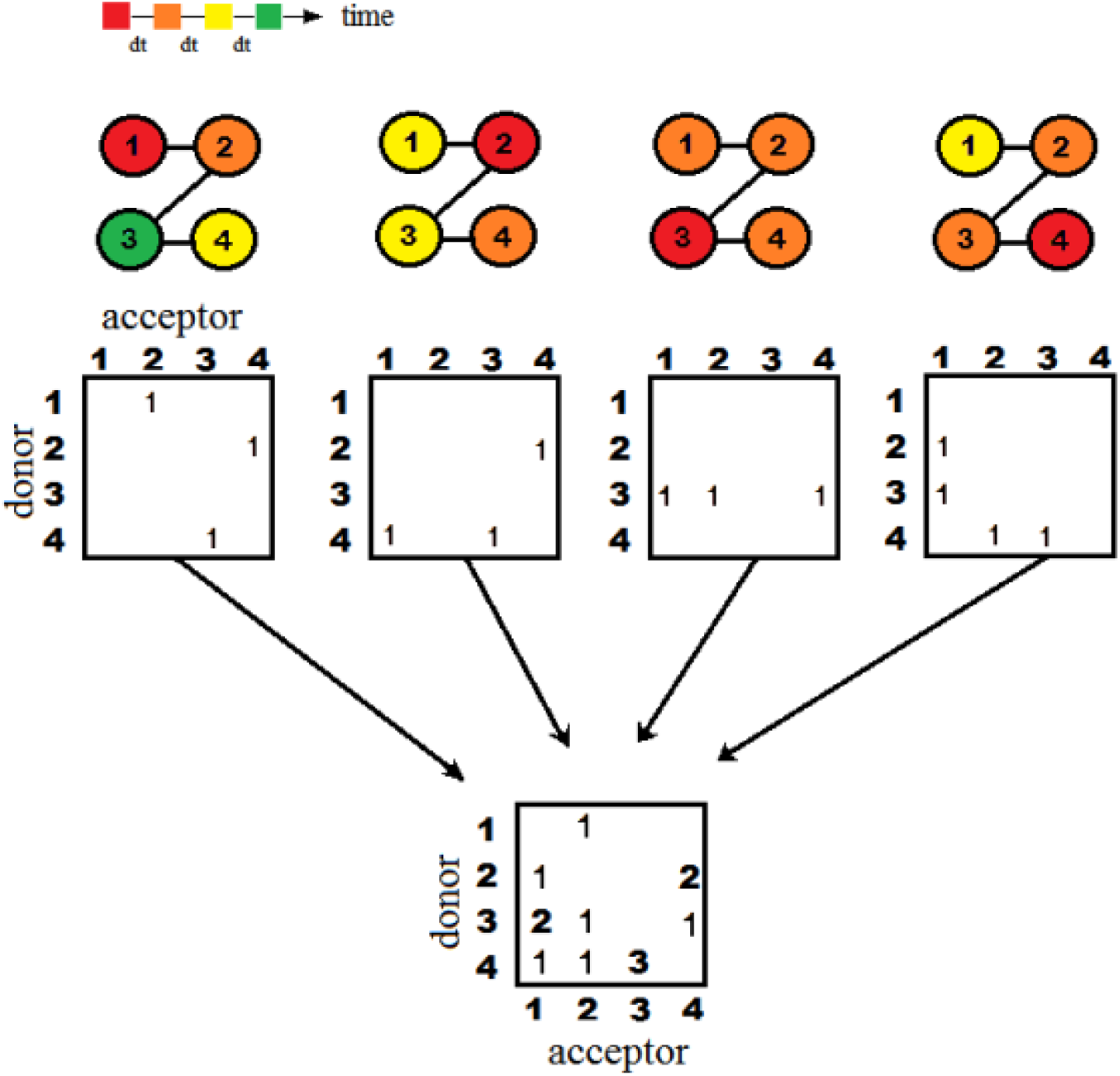
Recording the signal propagation into communication matrices. From the top, the number-labelled spheres linked by solid lines model a tetrapeptide (or four covalently bonded atoms), where the number is the residue (or atom) index. The colors indicate a time sequence of characteristic times (*t*c). The signals start from the red nodes and then propagate to orange, yellow and eventually green nodes in equal intervals. Donor-acceptor matrices are used to note the pairwise signal transmissions from a donor to an acceptor, if their *t*c differs by a selected time interval, dt = 0.2 ps in this study. Counts are added to a cell in the communication matrix corresponding to a given “connected” pair when perturbations are initiated from a number of nodes and in 21 different directions. Following eq. (3), after summation over all matrices with perturbation initiated at different node 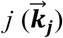, toward a selected set of directions, we can have the all-atom communication matrix *F*(*a*_*d*_, *a*_*r*_) that notes the residue pairs communicating the most.

**Figure 2.**
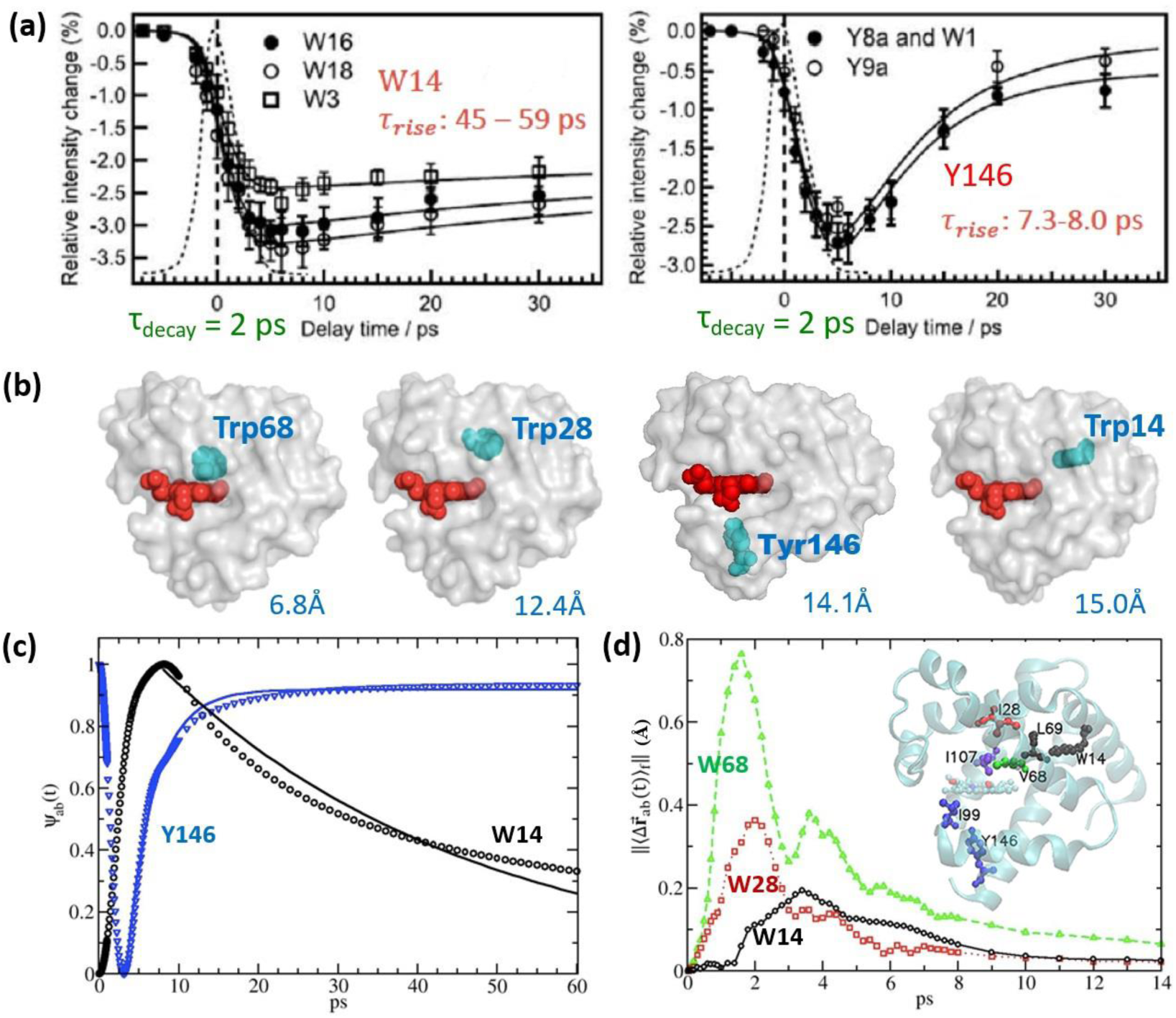
td-LRT using constant forces and impulse forces to explore biphasic relaxation dynamics in myoglobin. **(a)** and **(b)** are reproduced with minor modifications from the data reported in Sato et al[3] and Fujii et al.[2], respectively. **(a)** UV Resonance Raman (UVRR) data demonstrated the time responses of hydrophobic environment changes for W14 (left) and Y146 (right) upon photo-dissociation of monoxide (CO); the fast decay and slower recovery can be theoretically described by NMA-based CF-td-LRT[21]. In this case, CO association and dissociation caused permanent structural changes in myoglobin. **(b)** Without the binding or light-induced photo-dissociation of CO, heme was excited by a 405 nm pump pulse; the electronic relaxation of the heme induced specific modes of heme vibrations and the vibrational energy propagated from heme throughout the molecule. The signal is traced by pump-probed technique using Trp as the sensor to observe anti-stoke intensity changes. All the other tryptophan residues in the wild-type Mb were mutated into Phe or Tyr with aromatic side chains to maintain the structural integrity and the positions of interest were mutated into Trps to receive the measurable signals as a function of time. The distances between the heme (in red) and Trp sensors (V68W, I28W, Y146 and W14 colored in cyan; note that Y146 was not studied in Fujii et al.[2]) are denoted in the bottom right of the structures. Fujii et al. reported the time constant *τ*_*rise*_ of 3.0 ± 0.4 and 4.0 ± 0.6ps for for V68W and I28W, respectively, while W14 was not detectable.[2] **(c)** By using CF-td-LRT, the reaction coordinate *Ψ*_*ab*_(*t*), a normalized distance between the geometry centers of the residue pairs (the residues of interest studied by the UV-TR3 and the residues representing their hydrophobic environment) is plotted as a function of time. 350 PC modes up to ∼40 *cm*^−1^ (see Fig. 4) are used in the constant force td-LRT (CF-td-LRT) calculations. Data are fit to a single exponential function, A exp(−t/τ) or B − A exp(−t/τ) from t_0_ to 60 ps where t_0_ is at the end of the fast responses (where the spikes are seen) before the long-time relaxation. Relaxation time constants can be obtained from the fitting parameter, τ, revealing a relaxation of 39.0, and 3.5 ps for the residues W14, and Y146 that are characterized with a UVRR-observed relaxation of 4*9* ± 16 and 7.4 ± 1.4 ps, respectively. **(d)** The relative distance between center of mass of side chain, 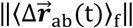, is investigated by the td-LRT using impulse forces (IF) (or IF-td-LRT), where the impulse forces include (1) a point force exerted at FE atoms pointed from CO (if bound) and (2) forces along the directions of the heme vibrational mode *ν*_7_.[8] The characteristic times of V68, I28 and W14 are 1.6, 2.0 and 3.4 ps, respectively. Our results agree with the time constants measured by UV-TR3 data, of which the time constants *τ*_*rise*_ for V68W and I28W are 3.0 ± 0.4 and 4.0 ± 0.6 ps, respectively. [2]

**Figure 3.**
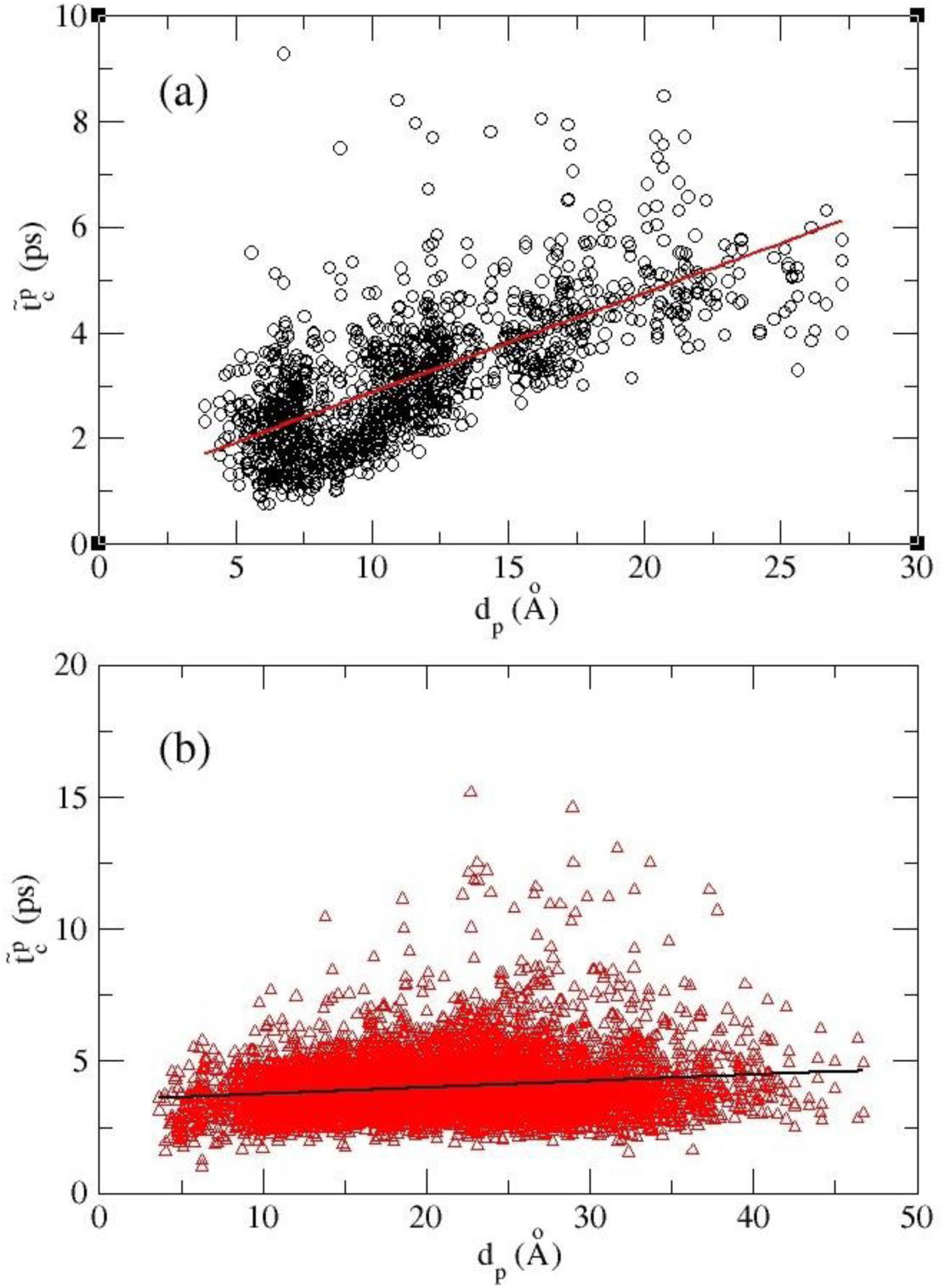
The site-to-site mean characteristic time, 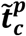, as a function of distances, *dp*, between pairs of perturbed sites and sensing sites. **(a)** The intra-helical signal propagation, where both perturbed and sensing sites are located at the same helix. **(b)** The inter-helical signal propagation, where perturbed and sensing sites are located at different helices. To estimate the intra-helical and inter-helical signal propagation speed, a linear regression is taken to give a speed (the inverse of the slopes) of 529 m/s for intra-helices **(a)** and that of 4076 m/s, a speed that is faster than sound propagation in water (1482 m/s at 20°C) [44] but slower than in steel (5930 m/s at 20°C),[45] for inter-helices. The correlation coefficient of 0.71 and 0.16 for (a) and (b), respectively, imply a more topology-dependent propagation in **(b)**.

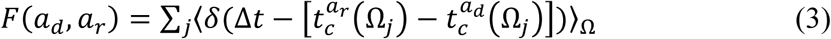

where *j* runs over all the selected *C*_α_ atoms being perturbed, δ(*x*) is a Kronecker delta function of *x*. Ω_*j*_ are forces orientation angles as defined in the previous section. The lower part of Figure 1 illustrates how communication matrix **F** is constructed by summing all the pairwise communications when signals are initiated from different sites in the system, one at a time. With the all-atom **F** matrix, we further define the residue level communication matrix (CM) by summing the counts belong to each residue pair, 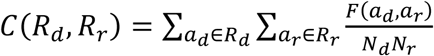, where *R*_*d*_ and *R*_*r*_ are donor and acceptor residues, respectively; *N*_*d*_ and *N*_*r*_ are the number of atoms in donor and acceptor residues, respectively.

To model the signals propagation process, we perturbed the residues within 6 Å from the ferrous ion (FE): F43, H64, V68, L89, H93, H97 and I99 represented in yellow spheres in the middle panel of Figure 5a. Consequently, 21 evenly distributed impulse forces are applied to each perturbation site and all the resulting signal communications were summed in a CM, *C*(*R*_*d*_, *R*_*r*_). In the CM, the diagonal elements are of intra-residue communication having high *C*(*R*_*d*_, *R*_*r*_) scores, which is intuitive but less meaningful in terms of the allostery. For the off-diagonal elements satisfying |*R*_*d*_ − *R*_*r*_| > 2, they provide information on the signal propagation between long range contacts (say, within or between secondary structures). Several hot spots that have high CM scores indicate their strategic locations to frequently communicate mechanical signals in multiple pathways. Then, a unique “communication score (CS)” can be assigned to each residue, which is defined as the highest CM score between this residue and any other non-neighboring residues in the protein – in other words, residue *i*’s CS is the highest score in either the *i*-th row or the *i*-th column (corresponding to donors or acceptors, respectively) of CM.

## Results

### Modeling the biphasic relaxation of residues of time-resolved ultrafast spectroscopy

It has been reported that the time constant measured by the resonance Raman spectra of tryptophan residues are sensitive to its hydrophobic environment change.[3, 41, 42] For the four measured residues in the UV-TR3 experiments: W14 on helix A,[2, 3] Y146 on helix H,[3] V68W on helix E,[2] and I28W on helix B,[2] their hydrophobic environment changes are monitored by the distance fluctuations between them and their closest hydrophobic residues (herein, distances for Trp14-Leu69, Ile28-I107, Val68-I107 and Tyr146-Ile99 pairs). The residues’ spatial distributions relative to heme can be found in Figure 2. To quantify the dynamics of corresponding residues, we define the reaction coordinate, 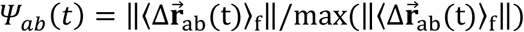, where 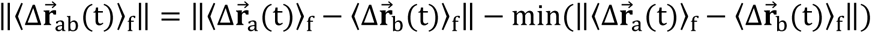 and 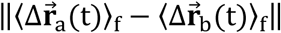 is the time-evolved distance between the side chain centers of residues a and b starting from when the force perturbations are introduced.

Considering conformational changes initiated by photodissociation of MbCO,[3] we use the CF-td-LRT particularly shown suitable for describing the conformational change.^[21]^ The time correlation functions (interchangeably used as time-dependent covariance) in eqn (1) are composed of 350 PC modes up to ∼40 *cm*^−1^ (see Figure 4). As described in the Methods, the constant forces that result from CO photodissociation can be derived from the known conformational changes and the inversion of the covariance matrix through time-independent linear response theory.^[21]^ With the derived forces and time-dependent covariance in eqn (1), the time-dependent conformational changes of Mb can be tracked. The normalized response curves *Ψ*_*ab*_(*t*) for the two pairs, W14-L69 and Y146-I99, can be drawn (Figure 2c). Fitting the data corresponding to the structural-change-associated slow relaxation (after the peak for W14 and after the minimum for Y146) to the exponential function A exp(−t/τ) or B − A exp(−t/ τ) reveals that the time constants τ obtained for the two pairs are respectively 39.0 and 3.5 ps, which are closely compared with the time constants 49.0 and 7.4 ps observed in time-resolved UVRR experiments.[3] It is worth noting that there is a fast rise or drop in *Ψ*_*ab*_(*t*) appearing in the first a few picoseconds before the signal partly (W14) or fully (Y146) recovers from a slower relaxation (Figure 2c). Revealed below, these fast responses can be described by td-LRT using impulse forces.

**Figure 4.**
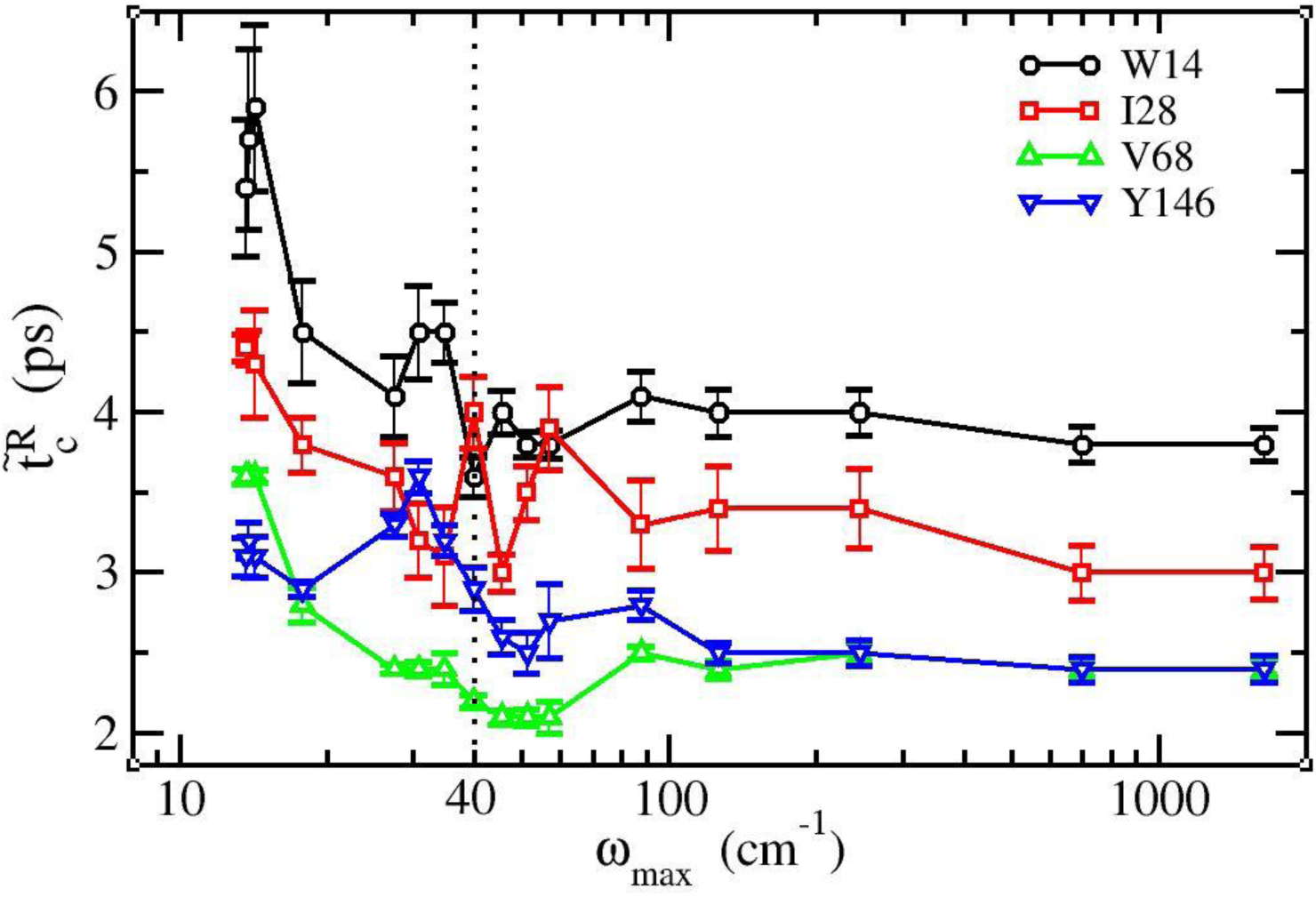
The characteristic time 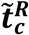 as a function of the cutoff frequency *ω*_*max*_ in the range of 12 < *ω*_*max*_ < 1100 *cm*^−1^ where ***ω***_***max***_ indicates the upper bound of the PC modes used in building the covariance matrix 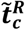 is the characteristic time over evenly distributed impulse forces acting on the FE atom only, and an error bar of 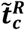 reflects the variance of characteristic times for constituent atoms in a residue. The data of the four residues W14, I28, V68 and Y146 studied by UVRR (see their distances from heme in Figure 2b) are shown in black circles, red squares, green upward-facing triangles and blue downward-facing triangles, respectively.

**Figure 5.**
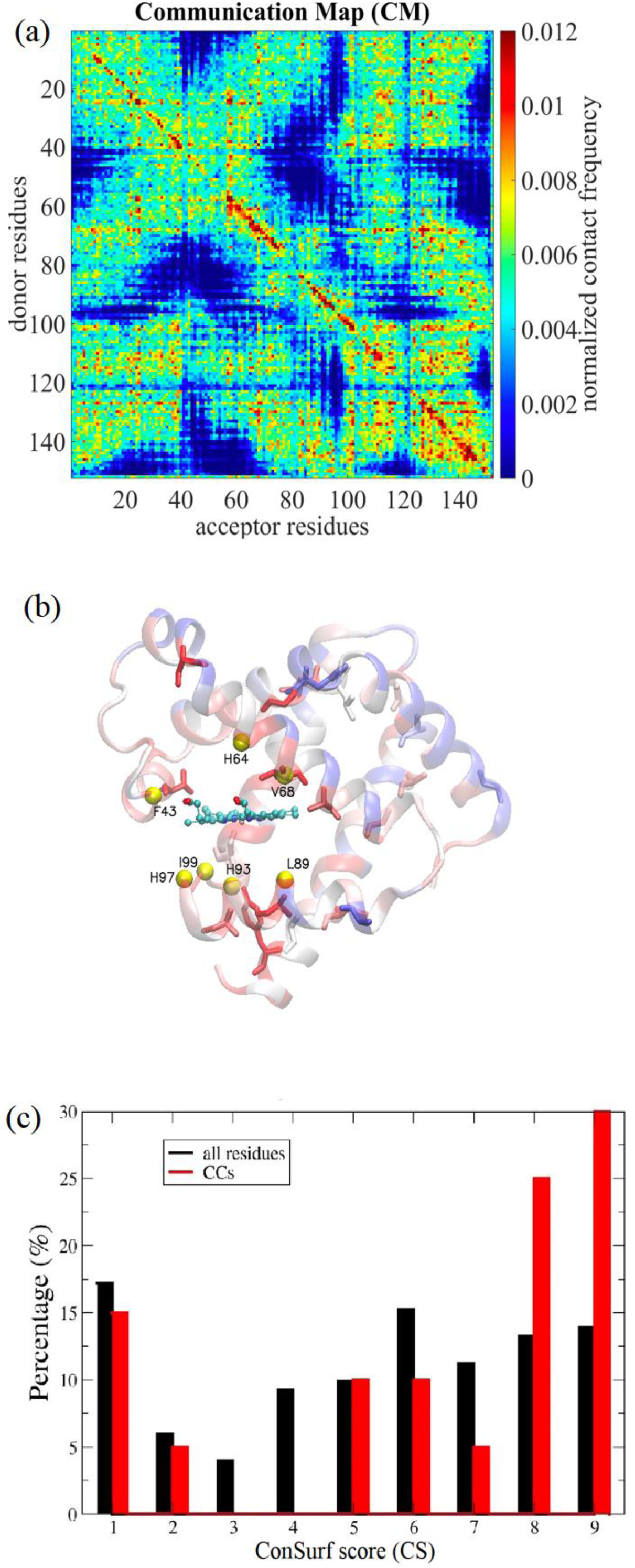
The analysis of communication centers (CCs). (top panel) The communication maps (CMs) are shown for Mb, respectively. **(middle panel)** The residues within 6 Å from FE - F43, H64, V68, L89, H93, H97 and I99 are chosen to be the perturbation sites, presented in yellow spheres. In CMs, the communication score of a residue is defined as the maximum the off-diagonal elements of the column (signal receiver) and row (signal donor) that contains the residue. Residues with the highest 20 communication scores are termed as the communication centers (CCs) of Mb, which are rank-ordered by their communication scores as V68, A134, A84, A130, G124, T39, G25, A71, A94, A90, A127, S58, H24, A143, G5, Y146, D141, K102, L115 and V114, presented in sticks; the colors of the ribbon diagram indicate residues’ evolutionary conservation from the least conserved (score 1, dark blue) to the most conserved (score 9, dark red) where the scores are derived from the multiple sequence alignment results provided by the Consurf server[46, 47]. The normalized histograms of the ConSurf scores from 1 to 9 of all the residues and CCs are colored in black and red for Mb **(lower panel)**. The result is consistent with our earlier data on dihydrofolate reductase where the CCs were shown to be evolutionarily conserved [17].

The relaxation time constants of another two tryptophan mutants V68 and I28 have been reported by the UV-TR3 experiment without significant conformational changes in Mb,[2] where the tryptophan band of anti-stokes intensities give the time constants for V68W and I28W, 3.0 ± 0.4 and 4.0 ± 0.6 ps, respectively.[2] We use the IF-td-LRT in eqn (2) to model the relaxation dynamics and the impulse force used to mimic the laser pump pulse is a force pointing from CO (if bound) to the FE atom perpendicular to heme plus the forces that model the “heme breathing” motion (*ν*_7_ mode).[8] Figure 2d shows the reaction coordinates, 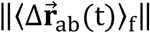, for residues V68, I28 and W14 labeled by green triangles, red squares and black circles, respectively. We define characteristic time *t*_*c*_ as the time when 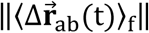 reaches its maximal amplitude. The *t*_*c*_ for 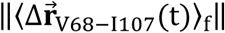 is 1.6 ps, shorter than 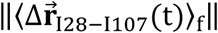 of 2.0 ps. The estimated fast responses are slightly faster but still close to the UV-TR3-observed time constants, 3.0 ± 0.4 and 4.0 ± 0.6 ps, respectively.[2] W14 has a *t*_*c*_ of 3.4 ps according to our IF-td-LRT while the signal is too weak to be detected in the UV-TR3 experiment.[2]

### Inter-helix signals propagate faster than intra-helix ones

The characteristic time *t*_*c*_ that indicates the arrival time of mechanical signals to individual atoms or residues enables us to examine how fast such signals propagate through secondary and tertiary protein structures. We quantify the signal propagation between (inter-) or within (intra-) α-helices, corresponding to perturbed sites and sensing sites located at different or the same helices, respectively.

It can be seen in Figure 3a that the intra-helical 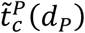(see symbol definition in Figure 3) increase with *d*_*P*_ linearly with a correlation coefficient of 0.71, where the linear regression gives a speed of 529 m/s, about twice slower than, 10 Å/ps (=1000 m/s)[43]. In comparison to the intra-helix signals, the 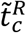 for signal propagation across different helices is much shorter than its intra-helical counterpart. The linear regression gives a speed of 4076 m/s, falling in between that in water (1482 m/s)[44] and in steel (5930 m/s),[45] for the inter-helical communication with a correlation coefficient of 0.16. The low linearity suggests a more topology-dependent nature of the inter-helical signal propagation than that for the intra-helical case. These results imply that the 3D packing of secondary structures can significantly accelerate the signal propagation speed, exemplified by the rigid molecule, myoglobin.

### The signal propagation mediated by selected modes

As a property derived from time-correlation functions used in IF-td-LRT, the characteristic time 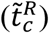 should be a function of constituent PC modes. To understand the mode dependence of characteristic time of residues of interest and therefore the inferred propagation, we calculate the 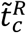 from the time dependent covariance matrix that comprises all the PC modes that are slower than a “cutoff frequency”, ω_*max*_ that goes from 12 to 1100 cm^-1^. Figure 4 shows the 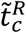 of the four residues examined in UV-TR3 experiments[2, 3] as a function of ω_*max*_ when the perturbation is introduced at the FE atom of the heme from 21 evenly distributed forces. When taking all the modes ≤1100 cm^-1^, we obtain 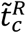 for I28, V68 and Y146 (note: this is the signal-residue response; not residue-environment responses as described before) as 3.0 ± 0.4, 2.4 ± 0.2 and 2.4 ± 0.3 ps, respectively, which are compatible with the UV-TR3 results 4.0 ± 0.6, 3.0 ± 0.4 and 2.0 ± 0.8 ps,[2, 3] respectively. However, we should note again that the experimental data for I28 and V68 were obtained from photoexcited Mb involving no conformational changes, while that for Y146 was measured from CO-photolyzed Mb with accompanied conformational changes, which involves multiple force perturbations on and near the heme. It can be seen in Fig4 (a) that 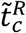 does not vary much until ω_*max*_ continues to drop below ∼40 *cm*^−1^ where monotonic increases of the 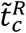 start to become apparent in all the four residues. Also, we found that PC modes larger than 100 *cm*^−1^ are barely influential to 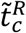. This could be because these modes are spatially localized and do not propagate the signals.

### The intramolecular communication centers are evolutionarily conserved

To investigate whether the residues frequently mediating the vibrational signals have any functional or evolutional importance, we define a set of residues as communication centers (CCs) (See Methods) which frequently communicate signals in multiple pathways. The CCs are functions of perturbing sites. Perturbing heme-binding residues located within 6 Å from the ferrous atom, F43, H64, V68, L89, H93, H97 and I99 (yellow balls in Fig. 5b), results in a communication map (CM) (Fig. 5a; see Methods). The CCs are the 20 residues having the highest communication scores (CSs; see Methods), which are rank-ordered as V68, A134, A84, A130, G124, T39, G25, A71, A94, A90, A127, S58, H24, A143, G5, Y146, D141, K102, L115 and V114 (Figure 4b). On the other hand, we calculate the evolutionarily conserved residues in Mb based on multiple sequence alignment results in the ConSurf database[46, 47] where residues are grouped into nine conservation levels by their “ConSurf scores” from 1 (most diverse) to 9 (most conserved) color-coded from blue to red in Figure 5b, respectively.

We then exam the distribution of the ConSurf scores for all the residues as well as for the CCs. It is found in the normalized histograms of CS (Figure 5c) that the average ConSurf score of CCs is 6.4, larger than the average ConSurf score for all the residues, 5.3. Also, there are 60% CCs of the ConSurf score value greater than or equal to 7 while only 38% of the residues in Mb meet the same criterion. Similarly, we previously found that the CCs are generally more conserved than average residues in dihydrofolate reductase (DHFR) that catalyzes the reduction of dihydrofolate (DHF) in the presence of the cofactor NADPH into terahydrofolate (THF) and NADP^+^. [17, 48, 49] These results suggest that CCs, bearing mechanical/communication importance, are evolutionarily conserved.

### Communication centers are not co-localized with functional mechanical hinges and folding cores with high local packing density

We further ask whether these conserved communication centers are a natural consequence of their important structural and mechanical roles in serving as folding cores[50] or being at the mechanical hinges.[50, 51] Among the 20 CCs in Mb, we found that there are seven folding cores (35% of CCs) including G25, A90, A94, V114, A127, A130 and Y146, which are identified by the fastest GNM mode peaks,[50] and the residue A71 is identified as a mechanical hinge by the slowest GNM mode (see Figure 6).[51, 52] However, all the other 12 CCs, including the top three communicators V68, A134, A84 (highlighted in Figure 6), are neither folding cores nor mechanical hinges, suggesting that these CCs are not properties readily derivable from proteins’ structural topology and low-frequency dynamics. Within the 12 CCs, T39, S58, V68, A134 and D141, having a ConSurf score ≥ 8, are highly conserved, while S58 (ConSurf score = 9), A134 and D141 are not anywhere close to the heme.

**Figure 6.**
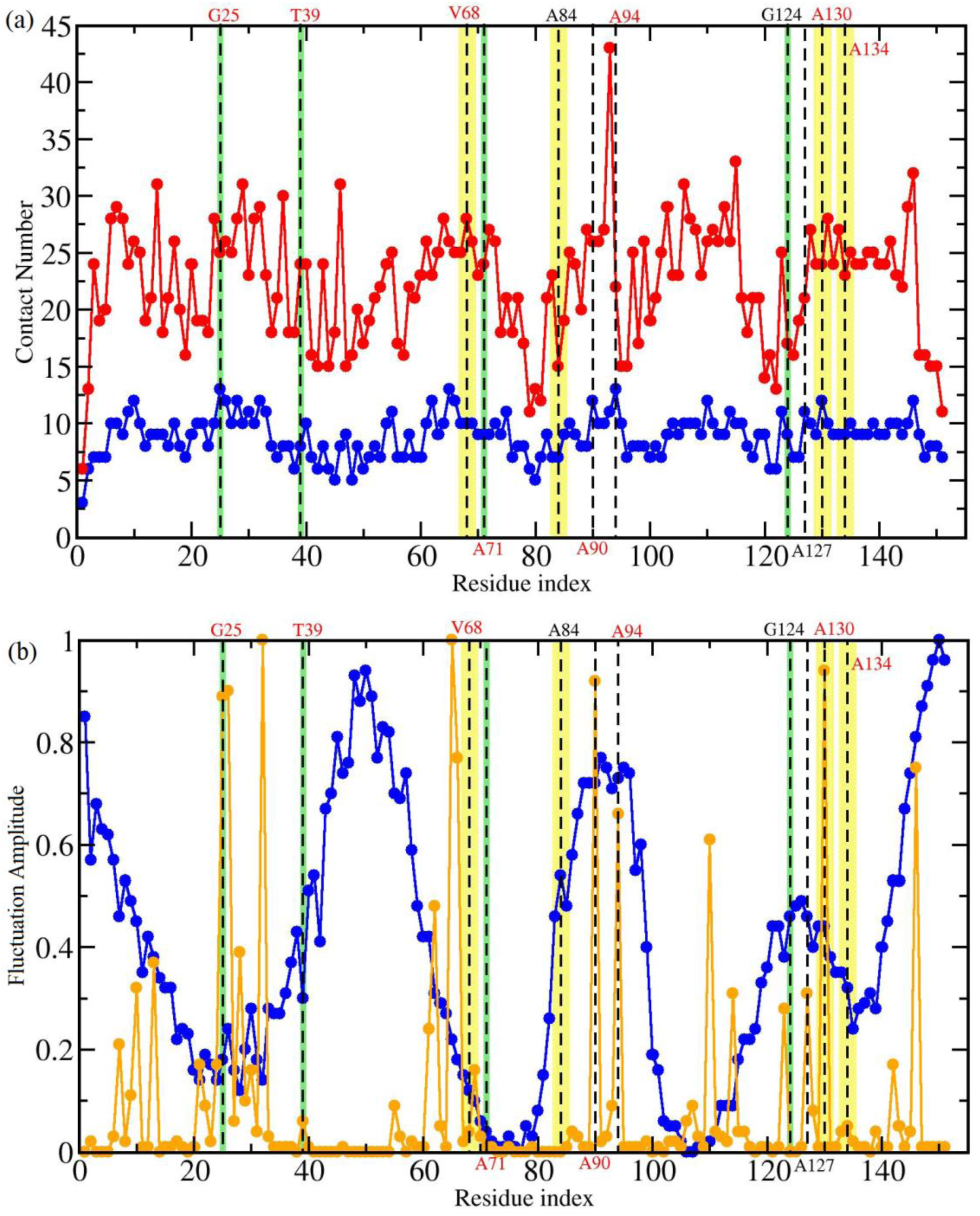
The contact number (a) and fluctuation amplitude (b) for Mb. Dash lines are on the residues that are deemed as the communication centers (CCs) derived by our IF-LRT. The top four communicators are highlighted in yellow; those that rank 5 to 8 are in green. In **(a)**, Per residue contact numbers based on atoms (red) and on residues (blue) are plotted along the residue index of Mb. In **(b)**, the frequency weighted fluctuation profiles comprising the slowest two modes (blue) and the fastest 10 modes (orange) are drawn. As one can clearly see in the figure, CCs are not necessarily located at highly packed/contacted regions, structural wise, neither are they co-localized with the slowest mode hinges [53-55] or the fastest mode peaks (indicating folding cores [50]), suggesting CC cannot be alternatively derived from apparent structural and dynamics properties.

## DISCUSSION

We have developed two td-LRTs, CF-tdLRT and IF-tdLRT, to model the long-time (involving conformational changes) and short-time relaxation (without conformational changes), respectively. Although the initial stage of the electronic excitation within the heme group involves quantum effects[5], the consequent relaxation process in protein environment seems qualitatively captured by our classical approach that describes the time characteristics of the energy transfer measured by the UVRR experiment. In this work, we also show that the PC modes can serve as a well-defined basis set to mediate the vibrational energy using the PCA-based tdLRTs. Three substantial issues are carefully investigated in this study: 1. How do we model the quantum external perturbation use a classical approach? 2. What kinds (frequency range of modes) of relaxation motions of the measured residues are captured by the time-dependent UVRR spectra? 3. Are there essential residues taking charge of the energy flow pathway?

In this study, the external perturbations are modeled by two kinds of classical forces: constant force induced by conformational changes and point impulse forces at heme. According to Mizutani and Kitagawa’s earlier study,[5] the laser pulse first excites the electronic state of FE atoms and then the excess heat relaxes to the vibrational state of heme before further propagating into the protein matrix. We describe the process of which by our IF-td-LRT using a point impulse force at FE atom along the direction perpendicular to the heme plus “effective forces” along the heme breathing (*ν*_7_) mode, observed by visible few-cycle-duration light pulses[8], due to aforementioned vibrational relaxation (Figure 2d and Figure 3). Second, the time constant of the measured residues could be attributed to the change of their hydrophobic environment.[3] In this work, we represent the hydrophobic environment of the measured residue using the displacement between the center of mass of the measured residues and its closest hydrophobic residue. From the Figure 2c and 2d, these approaches are shown to estimate the time constant appropriately. On the other hand, the corresponding band in UVRR spectrum for a measured residue could also be dominated by a specific set of vibrational modes,[2] and residues of interest itself instead of its hydrophobic environment can be monitored (Fig 4). In this aspect, we also measure the mean characteristic time of a residues,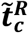, shown in Figure 4. For the full mode case, 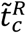 of I28, V68 and Y146 are 3.0 ± 0.4, 2.4 ± 0.2 and 2.4 ± 0.3 ps, respectively; it is compatible to the fast response results in UVRR experiment of 4.7 ± 1.3, 3.0 ± 0.7 and 2.0 ± 0.8 ps [2, 3], respectively. It has been shown in Figure 4 that the deletion of fast PC modes above 200 *cm*^−1^ only affect the time constants slightly, while the deletion of PC modes above ∼40 *cm*^−1^ cause the time constants fluctuate significantly. The results suggest that the medium frequency modes between 40 and 200 *cm*^−1^ play an important role in propagating signals. However it does not rule out the possible mechanism mediating the vibrational energy flow such like mode resonance or modes couplings [9, 24, 43].

Lastly, we address the third issue by building the time-dependent covariance matrix in eq.1 and eq.2 using selected modes. Launching signals from several active site residues, we can keep track of the accumulated count on the atom pairs where the signal flow through within a short time interval. Such a tally has shown a surprising accordance with remote residues’ importance in regulating protein function and their high conservation in evolution. The new “communication” property does not seem to co-localize with the mechanical hinges, previously shown to be functional relevant [36, 53, 54], or folding cores, separately defined by ENM’s slowest and the average of the fastest 10 modes (Fig 6). Similar phenomenon was observed in DHFR, where communication centers do not have to co-localize with high packing density or functional mechanical hinges [17].

## ACKNOWLEDGEMENTS

We acknowledge the computational resources supported by High Performance Computing Infrastructure (HPCI), Japan, and by National Center for High-performance Computing (NCHC) of National Applied Research Laboratories (NARLabs) of Taiwan. This work is funded by the Ministry of Science and Technology (104-2113-M-007-019, 106-2313-B-007-001- and 107-2313-B-007 -001 -) and National Center for Theoretical Sciences, Taiwan to L.W.Y.

## Supporting Materials

### Supporting Methods

#### A general time-independent linear response theory

Given the Hamiltonian of unperturbed system, *H*_0_, governs the dynamics of equilibrium system. The perturbed system is subject to force 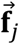 applied on atom *j*, and the Hamiltonian is 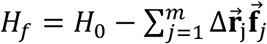, where 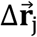 is the deviation from the mean of atom *j* due to the external force 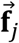. The time progression of the positional changes of atom *i* under external forces is of the form[21]

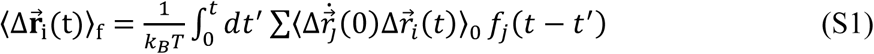

where *k*_*B*_ and *T* are the Boltzmann constant and temperature, respectively, 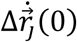 is the velocity of atom *j* at the moment when external forces, 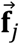, are applied. 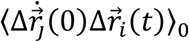 is the velocity-position time-correlation function sampled in the absence of perturbations noted by subscript 0, which can be expressed in the normal-mode space, where modes are treated as independent 1-D harmonic oscillators under solvent damping using the Langevin equation.[39] The detail derivation is referred to our previous work.[21]

#### The constant force time-dependent linear response theory (CF-td-LRT)

Substitute the force term in eq. (S1) with a time-invariant constant force 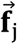, one can re-formulate eq. (S1) as

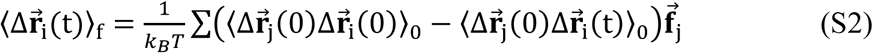

Let 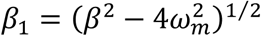 where *β* is the solvent friction and ω_*m*_ is the frequency of the PC mode *m*; for overdamped modes, 2ω_*m*_ < *β*, one can derive

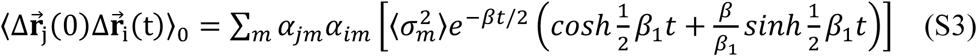

where *α*_*jm*_ and *α*_*im*_ are *j*-th and *i*-th component of the *m*-th PC mode, 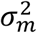 is the variance,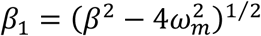; while an underdamped PC modes (2ω_*m*_ > *β*), one can have (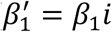 here)

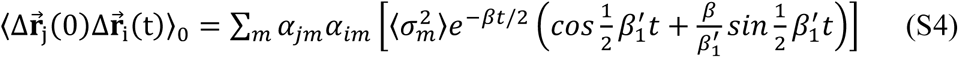

When time goes to infinite, the term 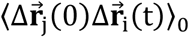 vanishes and eq. (S2) returns to the time independent form.[31]

#### The impulse force time-dependent linear response theory (IF-td-LRT)

Let the force in eq. (S1) be a delta function (the “impulse force”)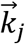, the format for the IF-td-LRT is

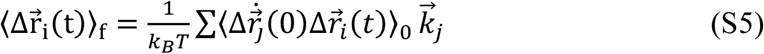

when 2ω_*m*_ < *β*

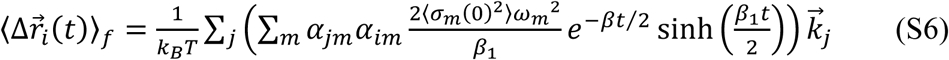

When 2ω_*m*_ > *β* (let 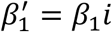; note that sinh *θ* = −*i* sin *iθ*),

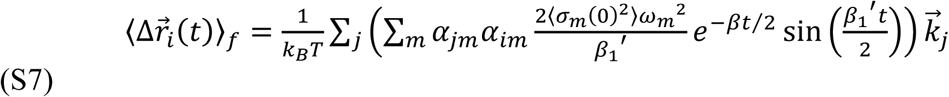

#### LRT for systems evolving on a “harmonic energy surface” can be directly derived without the need to assume small perturbations 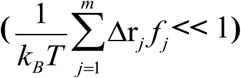

The validity of the aforementioned time-independent linear response theory (ti-LRT) equality[31] holds up on the fact that 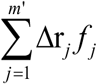 has to be enough small as compared to *kB*T (i.e. 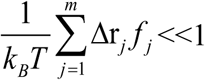). In general, this is valid if structural changes are small such as in the myoglobin case when the NMA treatment of the unperturbed covariance matrix is considered suitable[21]. However, when the induced structural changes are large,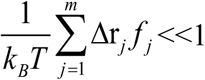 may not hold. However, we show below how the theory still holds when the protein conformational changes are not negligibly small. It serves the foundation for us to apply the theory to MD-derived trajectory subject to quasi-harmonic analyses.

Unperturbed Hamiltonian Ξ_0_, though not necessarily in the local minima, can be approximated harmonically from a classic-forcefield-defined or elastic-potential-defined energy minima such that

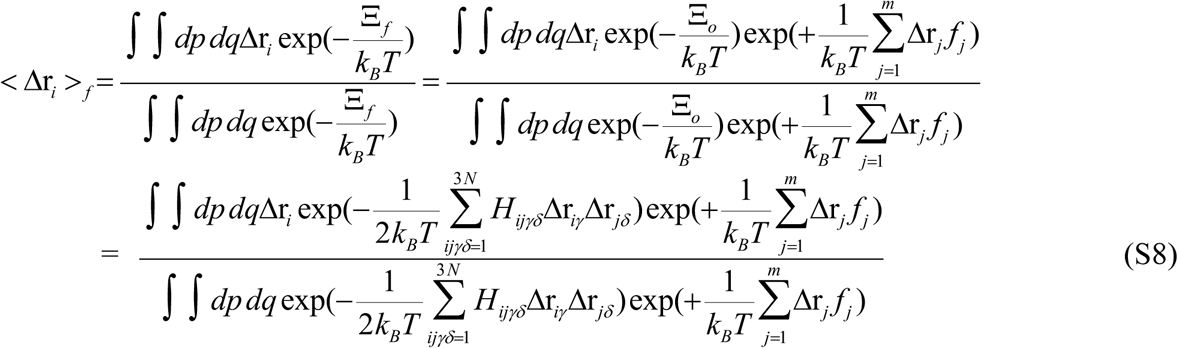

Let 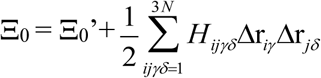; Ξ_0_’ is the energy minimum about which the harmonic approximation of the potential takes place. *H*_*ijγδ*_ are the components of Hessian (force constant matrix) **H**; *γ, δ* denote the Cartesian x, y and z. The constant term 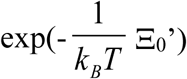 vanish as being factored out from both the numerator and the denominator. Assuming fj is readily available (possibly from the standard version of ti-LRT), to avoid the restriction that 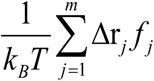 has to be small at the standard LRT case, we would like to directly integrate the above equation (now rewritten in its matrix-vectorial form) such that

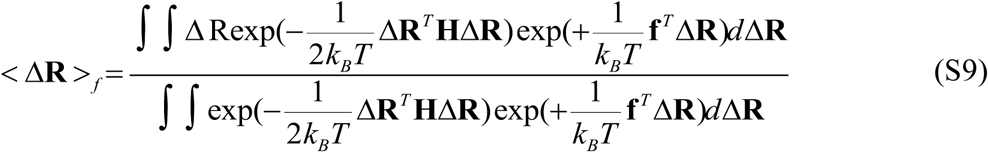

where conformational changes Δ**R** and forces **f** are 3*N*-d column vectors;

For the definite integral in the numerator and denominator, the equality

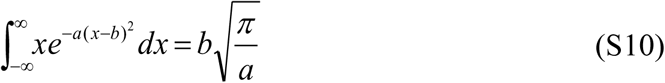

and

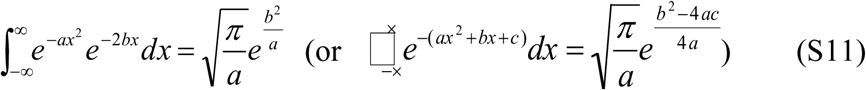

can be used respectively.

Using Eq. S10,

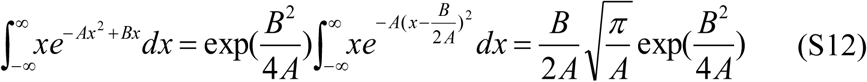

Using Eq. S11,

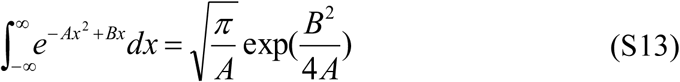

(S9) is of the ratio (S12) to (S13), which is 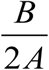, where A can be viewed as 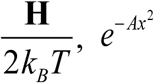 as 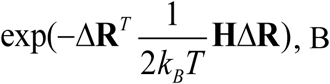, B as 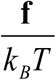 and *e*^*Bx*^ as exp(+*²***f** ^*T*^ Δ**R**).

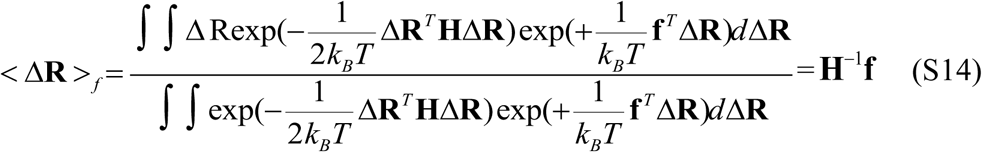

According to the NMA theories[50, 51], 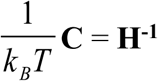(where elements in **C** are C_ij*γδ*_= < Δr_*iγ*_ Δr_*jδ*_ >) if the protein’s Hamiltonian is at the energy minimum and its adjacent potential surface is harmonically approximated.

Hence,

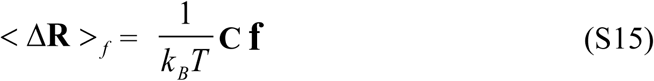

which arrives the same formula as shown before.[21, 29-31]

Therefore, when 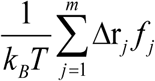 is not necessarily small (e.g. the ligand (or protein)-induced conformational changes that can be seen in long MD simulation or coarse-grained models), the same equation can be used if its potential is harmonically approximated.

## References

[1] Lim, M., Jackson, T.A. & Anfinrud, P.A. Femtosecond Near-IR Absorbance Study of Photoexcited Myoglobin: Dynamics of Electronic and Thermal Relaxation. The Journal of Physical Chemistry 100, 12043–12051 (1996).

[2] Fujii, N., Mizuno, M., Ishikawa, H. & Mizutani, Y. Observing Vibrational Energy Flow in a Protein with the Spatial Resolution of a Single Amino Acid Residue. The Journal of Physical Chemistry Letters 5, 3269–3273 (2014).

[3] Sato, A., Gao, Y., Kitagawa, T. & Mizutani, Y. Primary protein response after ligand photodissociation in carbonmonoxy myoglobin. Proceedings of the National Academy of Sciences of the United States of America 104, 9627–9632 (2007).

[4] Sagnella, D.E., Straub, J.E., Jackson, T.A., Lim, M. & Anfinrud, P.A. Vibrational population relaxation of carbon monoxide in the heme pocket of photolyzed carbonmonoxy myoglobin: comparison of time-resolved mid-IR absorbance experiments and molecular dynamics simulations. Proceedings of the National Academy of Sciences of the United States of America 96, 14324–14329 (1999).

[5] Mizutani, Y. Direct Observation of Cooling of Heme Upon Photodissociation of Carbonmonoxy Myoglobin. Science 278, 443–446 (1997).

[6] Champion, P.M. Chemistry. Following the flow of energy in biomolecules. Science 310, 980–982 (2005).

[7] Sage, J.T., Durbin, S.M., Sturhahn, W., Wharton, D.C., Champion, P.M., Hession, P., et al. Long-Range Reactive Dynamics in Myoglobin. Physical Review Letters 86, 4966–4969 (2001).

[8] Armstrong, M.R., Ogilvie, J.P., Cowan, M.L., Nagy, A.M. & Miller, R.J. Observation of the cascaded atomic-to-global length scales driving protein motion. Proceedings of the National Academy of Sciences of the United States of America 100, 4990–4994 (2003).

[9] Gao, Y., El-Mashtoly, S.F., Pal, B., Hayashi, T., Harada, K. & Kitagawa, T. Pathway of information transmission from heme to protein upon ligand binding/dissociation in myoglobin revealed by UV resonance raman spectroscopy. J Biol Chem 281, 24637–24646 (2006).

[10] Kondoh, M., Mizuno, M. & Mizutani, Y. Importance of Atomic Contacts in Vibrational Energy Flow in Proteins. J Phys Chem Lett 7, 1950–1954 (2016).

[11] Levantino, M., Schiro, G., Lemke, H.T., Cottone, G., Glownia, J.M., Zhu, D., et al. Ultrafast myoglobin structural dynamics observed with an X-ray free-electron laser. Nat Commun 6, 6772 (2015).

[12] Sharp, K. & Skinner, J.J. Pump-probe molecular dynamics as a tool for studying protein motion and long range coupling. Proteins 65, 347–361 (2006).

[13] Ghosh, A. & Vishveshwara, S. A study of communication pathways in methionyl-tRNA synthetase by molecular dynamics simulations and structure network analysis. Proceedings of the National Academy of Sciences of the United States of America 104, 15711–15716 (2007).

[14] Chennubhotla, C. & Bahar, I. Signal propagation in proteins and relation to equilibrium fluctuations. PLoS computational biology 3, 1716–1726 (2007).

[15] Chennubhotla, C. & Bahar, I. Markov propagation of allosteric effects in biomolecular systems: application to GroEL-GroES. Molecular systems biology 2, 36 (2006).

[16] Ota, N. & Agard, D.A. Intramolecular signaling pathways revealed by modeling anisotropic thermal diffusion. Journal of molecular biology 351, 345–354 (2005).

[17] Huang, B.-C., Chang-Chein, C.-H. & Yang, L.-W. Intramolecular Communication and Allosteric Sites in Enzymes Unraveled by Time-Dependent Linear Response Theory. https://www.biorxiv.org/content/10.1101/677617v1; submitted for peer review, (2019).

[18] Chen, J., Dima, R.I. & Thirumalai, D. Allosteric communication in dihydrofolate reductase: signaling network and pathways for closed to occluded transition and back. Journal of molecular biology 374, 250–266 (2007).

[19] Sagnella, D.E., Straub, J.E. & Thirumalai, D. Time scales and pathways for kinetic energy relaxation in solvated proteins: Application to carbonmonoxy myoglobin. The Journal of Chemical Physics 113, 7702 (2000).

[20] Hayward, S., Kitao, A., Hirata, F. & Go, N. Effect of solvent on collective motions in globular protein. Journal of molecular biology 234, 1207–1217 (1993).

[21] Yang, L.W., Kitao, A., Huang, B.C. & Go, N. Ligand-induced protein responses and mechanical signal propagation described by linear response theories. Biophysical journal 107, 1415–1425 (2014).

[22] Leitner, D.M. Frequency-resolved communication maps for proteins and other nanoscale materials. J Chem Phys 130, 195101 (2009).

[23] Gnanasekaran, R., Agbo, J.K. & Leitner, D.M. Communication maps computed for homodimeric hemoglobin: computational study of water-mediated energy transport in proteins. J Chem Phys 135, 065103 (2011).

[24] Yu, X. & Leitner, D.M. Heat flow in proteins: computation of thermal transport coefficients. J Chem Phys 122, 54902 (2005).

[25] Moritsugu, K., Miyashita, O. & Kidera, A. Vibrational Energy Transfer in a Protein Molecule. Physical Review Letters 85, 3970–3973 (2000).

[26] Henry, E.R., Eaton, W.A. & Hochstrasser, R.M. Molecular dynamics simulations of cooling in laser-excited heme proteins. Proceedings of the National Academy of Sciences of the United States of America 83, 8982–8986 (1986).

[27] Zhang, Y., Fujisaki, H. & Straub, J.E. Molecular dynamics study on the solvent dependent heme cooling following ligand photolysis in carbonmonoxy myoglobin. J Phys Chem B 111, 3243–3250 (2007).

[28] Sagnella, D.E. & Straub, J.E. Directed Energy “Funneling” Mechanism for Heme Cooling Following Ligand Photolysis or Direct Excitation in Solvated Carbonmonoxy Myoglobin. The Journal of Physical Chemistry B 105, 7057–7063 (2001).

[29] Essiz, S.G. & Coalson, R.D. Dynamic linear response theory for conformational relaxation of proteins. J Phys Chem B 113, 10859–10869 (2009).

[30] Kubo, R. The fluctuation-dissipation theory. Report Progress Physics 29, (1966).

[31] Ikeguchi, M., Ueno, J., Sato, M. & Kidera, A. Protein Structural Change Upon Ligand Binding: Linear Response Theory. Physical Review Letters 94, 078102 (2005).

[32] Kitao, A., Hayward, S. & Go, N. Energy landscape of a native protein: Jumpingamong-minima model. Proteins: Structure, Function, and Genetics 33, 496–517 (1998).

[33] Kitao, A. & Go, N. Investigating protein dynamics in collective coordinate space. Current Opinion in Structural Biology 9, 164–169 (1999).

[34] Chang, K.C., Salawu, E.O., Chang, Y.Y., Wen, J.D. & Yang, L.W. Resolution-exchanged structural modeling and simulations jointly unravel that subunit rolling underlies the mechanism of programmed ribosomal frameshifting. Bioinformatics 35, 945–952 (2019).

[35] Phillips, J.C., Braun, R., Wang, W., Gumbart, J., Tajkhorshid, E., Villa, E., et al. Scalable molecular dynamics with NAMD. Journal of computational chemistry 26, 1781–1802 (2005).

[36] Yang, L.W., Eyal, E., Bahar, I. & Kitao, A. Principal component analysis of native ensembles of biomolecular structures (PCA_NEST): insights into functional dynamics. Bioinformatics 25, 606–614 (2009).

[37] Kabsch, W. A solution for the best rotation to relate two sets of vectors. Acta Crystallographica Section A 32, 922–923 (1976).

[38] Kitao, A., Hirata, F. & Gō, N. The effects of solvent on the conformation and the collective motions of protein: Normal mode analysis and molecular dynamics simulations of melittin in water and in vacuum. Chemical Physics 158, 447–472 (1991).

[39] Chandrasekhar, S. Stochastic Problems in Physics and Astronomy. Reviews of Modern Physics 15, 1–89 (1943).

[40] Acbas, G., Niessen, K.A., Snell, E.H. & Markelz, A.G. Optical measurements of long-range protein vibrations. Nat Commun 5, 3076 (2014).

[41] Efremov, R.G., Feofanov, A.V. & Nabiev, I.R. Effect of hydrophobic environment on the resonance Raman spectra of tryptophan residues in proteins. Journal of Raman Spectroscopy 23, 69–73 (1992).

[42] Chi, Z. & Asher, S.A. UV Raman Determination of the Environment and Solvent Exposure of Tyr and Trp Residues. The Journal of Physical Chemistry B 102, 9595–9602 (1998).

[43] Leitner, D.M. Vibrational Energy Transfer in Helices. Physical Review Letters 87, 188102 (2001).

[44] Haynes, W.M. CRC Handbook of Chemistry and Physics. CRC Press, (2012).

[45] Dr. rer. nat. Dr.-Ing. E. h. Josef Krautkrämer, D.r.n.H.K. Ultrasonic Testing of Materials. Springer Berlin Heidelberg, (1990).

[46] Landau, M., Mayrose, I., Rosenberg, Y., Glaser, F., Martz, E., Pupko, T., et al. ConSurf 2005: the projection of evolutionary conservation scores of residues on protein structures. Nucleic acids research 33, W299–302 (2005).

[47] Ashkenazy, H., Erez, E., Martz, E., Pupko, T. & Ben-Tal, N. ConSurf 2010: calculating evolutionary conservation in sequence and structure of proteins and nucleic acids. Nucleic acids research 38, W529–533 (2010).

[48] Fierke, C.A., Johnson, K.A. & Benkovic, S.J. Construction and evaluation of the kinetic scheme associated with dihydrofolate reductase from Escherichia coli. Biochemistry 26, 4085–4092 (1987).

[49] Boehr, D.D., McElheny, D., Dyson, H.J. & Wright, P.E. The dynamic energy landscape of dihydrofolate reductase catalysis. Science 313, 1638–1642 (2006).

[50] Bahar, I., Atilgan, A.R., Demirel, M.C. & Erman, B. Vibrational Dynamics of Folded Proteins: Significance of Slow and Fast Motions in Relation to Function and Stability. Physical Review Letters 80, 2733–2736 (1998).

[51] Yang, L.W., Liu, X., Jursa, C.J., Holliman, M., Rader, A.J., Karimi, H.A., et al. iGNM: a database of protein functional motions based on Gaussian Network Model. Bioinformatics 21, 2978–2987 (2005).

[52] Li, H., Chang, Y.Y., Yang, L.W. & Bahar, I. iGNM 2.0: the Gaussian network model database for biomolecular structural dynamics. Nucleic acids research 44, D415–422 (2016).

[53] Yang, L.W. & Bahar, I. Coupling between catalytic site and collective dynamics: a requirement for mechanochemical activity of enzymes. Structure 13, 893–904 (2005).

[54] Chandrasekaran, A., Chan, J., Lim, C. & Yang, L.W. Protein Dynamics and Contact Topology Reveal Protein-DNA Binding Orientation. J Chem Theory Comput 12, 5269–5277 (2016).

[55] Li, H., Sakuraba, S., Chandrasekaran, A. & Yang, L.W. Molecular binding sites are located near the interface of intrinsic dynamics domains (IDDs). J Chem Inf Model 54, 2275–2285 (2014).

